# Cross-task contributions of fronto-basal ganglia circuitry in response inhibition and conflict-induced slowing

**DOI:** 10.1101/199299

**Authors:** Sara Jahfari, K Richard Ridderinkhof, Anne GE Collins, Tomas Knapen, Lourens J Waldorp, Michael J Frank

**Author notes:** **Corresponding Author:** Sara Jahfari, Department of Cognitive Psychology, Vrije Universiteit van Amsterdam, Van der Boechorststraat 1, 1018 BT Amsterdam, The Netherlands, Or, Michael J. Frank, Department of Cognitive, Linguistic and Psychological Sciences and Brown Institute for Brain Sciences, Brown University, 190 Thayler St, Providence RI 02912-1821, USA.

## Abstract

Why are we so slow in choosing the lesser of two evils? We considered whether such slowing relates to uncertainty about the value of these options, which arises from the tendency to avoid them during learning, and whether such slowing relates to fronto-subthalamic inhibitory control mechanisms. 49 participants performed a reinforcement-learning task and a stop-signal task while fMRI was recorded. A reinforcement-learning model was used to quantify learning strategies. Individual differences in lose-lose slowing related to information uncertainty due to sampling, and independently, to less efficient response inhibition in the stop-signal task. Neuroimaging analysis revealed an analogous dissociation: subthalamic nucleus (STN) BOLD activity related to variability in stopping latencies, whereas weaker fronto-subthalamic connectivity related to slowing and information sampling. Across tasks, fast inhibitors increased STN activity for successfully cancelled responses in the stop task, but decreased activity for lose-lose choices. These data support the notion that fronto-STN communication implements a rapid but transient brake on response execution, and that slowing due to decision uncertainty could result from an inefficient release of this “hold your horses” mechanism.

## Introduction

Optimal foraging entails learning to select among decision alternatives, based on their (hidden) probabilistic values. Individuals differ in their exploration/exploitation balance, and hence the degree to which they sample options with lower valued outcomes, during reinforcement learning. Such inter-individual variability may help understand the mechanisms involved in choosing the lesser of two evils. Value-based decision-making often requires choice between options that have similar learned values but may never have been presented together (for example, a novel choice between miso soup and corn chowder). These kinds of choices can elicit conflict arising from either the novel pairing of two previously desired outcomes (win-win) or undesired outcomes (lose-lose). Despite identical value differences, the novel pairing of two lose-lose options is consistently associated with prolonged decision times when compared to win-win conflict^1–5^. While the relative speeding for high valued options is attributed to effects of reward expectation (and dopamine levels) on reaction time (RT), the literature has generally not considered the impact of differential uncertainty about choice values. Consider a common reinforcement-learning task in which an agent learns to choose among pairs of options with different reinforcement probabilities (e.g., 80% vs. 20%, 70% vs. 30%, and 60% vs. 40%)^6^. While one can optimize rewards in this task by exploiting/maximizing (i.e., always choosing the more rewarded option), this strategy would prevent the agent from exploration and hence from acquiring a precise representation about the value of the lesser options^7,8^. Critically, this exploitation strategy would also then make it more difficult to later choose between a 40% and 20% option (a high-conflict lose-lose choice), due to less sampling and greater uncertainty about their true values.

What are the neural mechanisms that can leverage such uncertainty to adjust decision times? Prior studies indicate that when presented with decision conflict, increased activity in the STN acts to delay response execution by inhibiting action altogether^9–11^ or by raising the decision threshold, i.e., the level of evidence required to make a choice^1,2,4,12–18^. Intuitively, a common mechanism for response inhibition and threshold adjustment seemingly implies that faster or more efficient inhibition would relate to more conflict-induced slowing. However, in the case of lose-lose conflict such a fixed increase in decision threshold mechanism is maladaptive when the learned information for the optimal choice is sparse (i.e., it could engender decision paralysis). Instead, simulation studies suggested that the STN “hold your horses” mechanism is dynamic, with a fast initial STN surge that is followed by a steep decline of activation (“releasing the horses”), facilitating choice even when the evidence is sparse^4,15^. This dynamic could even suggest an efficient initial STN surge, and hence rapid response inhibition, might actually lead to *less* uncertainty-induced slowing.

We aimed to specify these relationships with the examination of two tasks. Functional magnetic resonance imaging (fMRI) data was recorded while participants performed a reinforcement-learning task followed by a test-phase containing novel win-win, lose-lose and win-lose pairs without feedback (Fig. 1a). Here, the degree of exploration/exploitation (and hence subsequent uncertainty in learned values of lose options) was assessed during learning by stochasticity in choices and quantified with a reinforcement-learning model that reliably predicted trial-to-trial choices (Fig. 2). Importantly, we also administered a stop-signal task to assess the efficiency of response inhibition in the absence of learning (Fig. 1b), and to relate this to the behavioral and neural markers of conflict-based slowing. We assess how choice strategies and the efficiency of response inhibition each relate to slowing in 1) reaction times, 2) the BOLD response of the STN, and 3) the strength of effective connectivity in the fronto-subthalamic pathway by using a model-driven effective connectivity approach termed ancestral graphs^19^. This last explorative analysis followed prior studies suggesting that the communication from PFC into the STN (the so-called hyperdirect pathway^20^), is enhanced under response conflict^10,13,21,22^ to motivate a brake^23–25^, or decision threshold adjustments on striatal reward-based choice in order to prevent impulsive or premature responses^2,12,15,26^.

**Figure 1.**
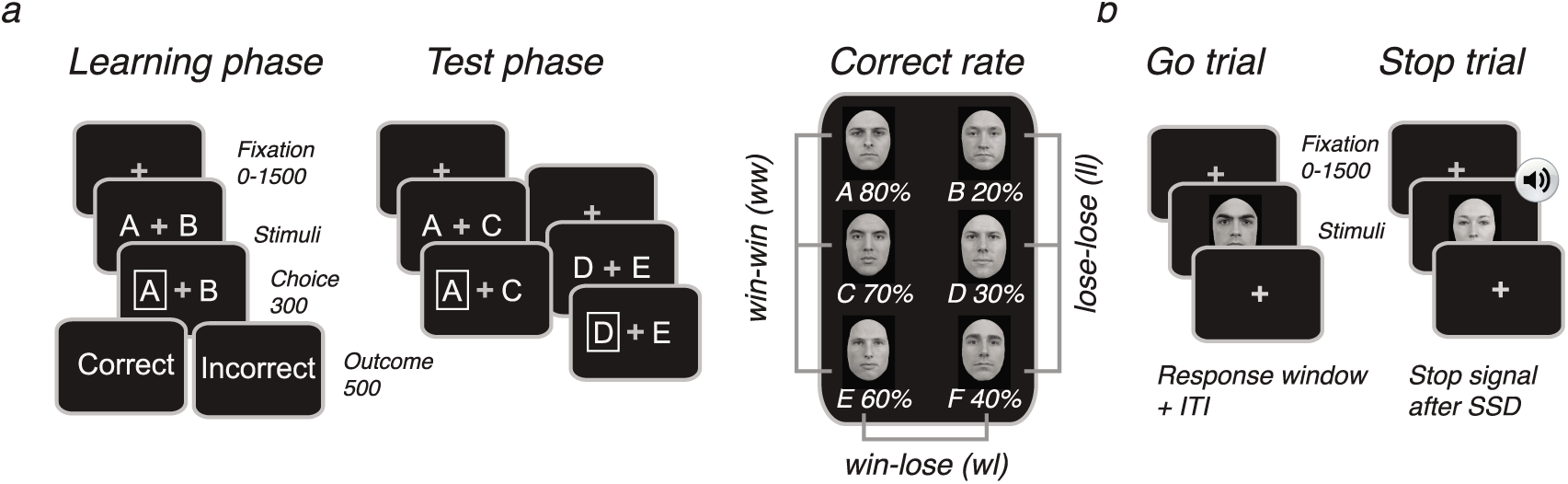
Experimental design. a) Reinforcement-learning task. During learning, two faces were presented at each trial, and participants learned to select the most optimal face stimulus (A, C, E) solely through probabilistic feedback (probability of correct is displayed beneath each stimulus). The learning-phase only contained three face pairs (AB, CD, ED) for which feedback was given. In the test-phase, faces were arranged into 15 combinations. Trials were further identical to the learning-phase with the exception of feedback. b) Stop-signal task. Each trial started with the presentation of a fixation-cross followed by a male or female face stimulus, indicating a left or right response. During stop trials, a tone was played at a variable delay (SSD) after the presentation of the go stimulus. The tone instructed participants to suppress the indicated response.

**Figure 2.**
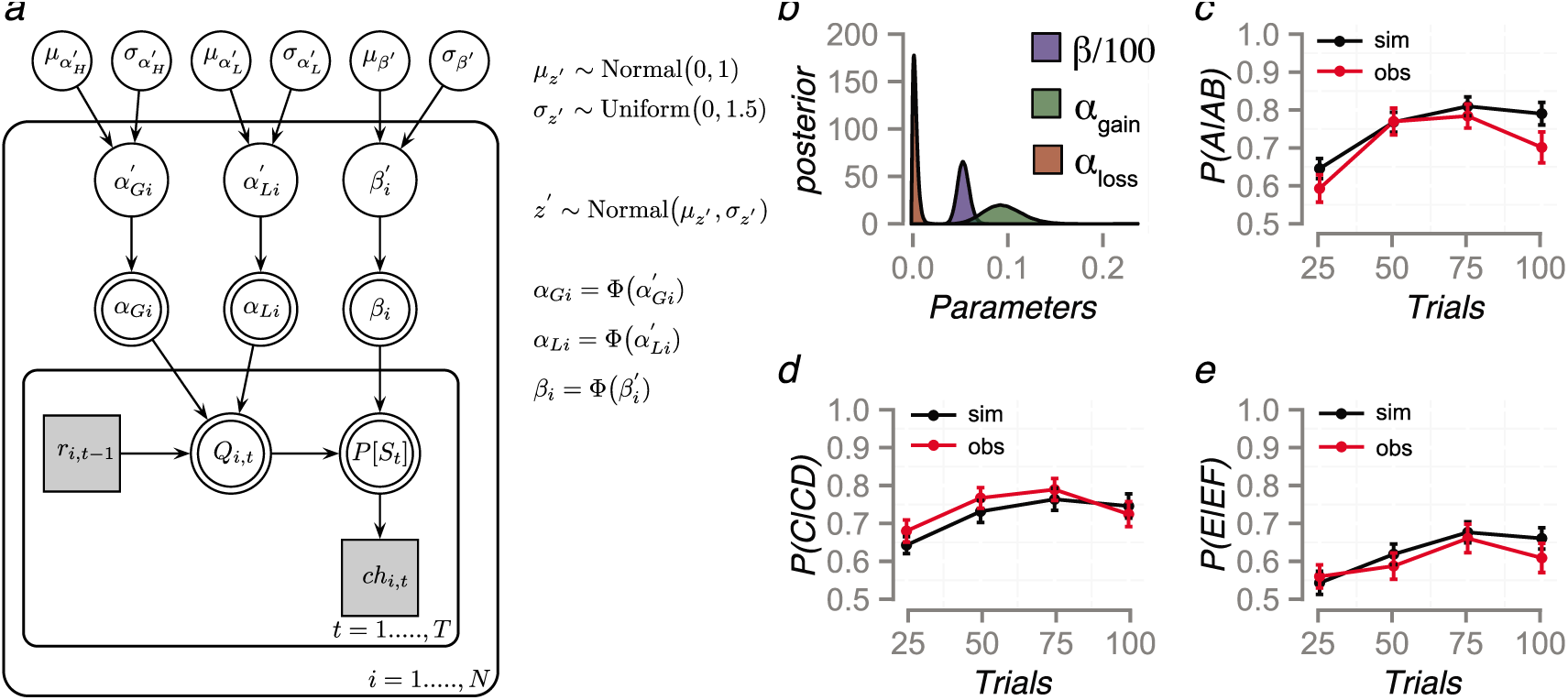
Q-learning model and performance. Graphical Q-learning model for hierarchical Bayesian parameter estimation (a). Փ () is the cumulative standard normal distribution function. The model consists of an outer subject (i=1,…..,N), and an inner trial plane (t=1,…,T). Nodes represent variables of interest. Arrows are used to indicate dependencies between variables. Double borders indicate deterministic variables. Continues variables are denoted with circular nodes, and discrete with square nodes. Observed variables are shaded in grey. The right panel shows group-level posteriors for all Q-learning parameters (with β/100) (b), and model performance where data is simulated with the estimated parameters and evaluated against the observed data for the AB (c), CD (d), or EF (e) pairs. Error bars represent SEM.

## RESULTS

49 young adults (25 male; mean age = 22 years; range 19-29 years) participated in this study. Four participants were excluded from all analyses due to movement (2), incomplete sessions (1), or misunderstanding of task instructions (1). One participant did not complete the stop-task, and for one we were unable to obtain reliable SSRT estimates (stopping latency) therefore they were only included for the RL-task analysis. As shown in Fig. 1, participants performed a reinforcement-learning task^27^ and a stop-signal task in the MRI scanner. In our reinforcement learning task participants learned to select among choices with different probabilities of reinforcement (i.e., AB 80:20, CD 70:30, and EF 60:40). A subsequent test-phase, where feedback was omitted, required participants to select the optimal option among novel pairs involving low (win-lose) or high (win-win and lose-lose) decision conflict.

### Uncertainty and Conflict-induced slowing

During the test-phase, as expected, accuracy in choosing the more rewarded stimulus was reduced for both high conflict lose-lose and win-win pairs compared to low-conflict win-lose pairs (F(2,88)=18.8, p<0.0001; Fig. 3a). Slowed RT’s were observed for only the lose-lose pairs (F(2,88)=21.4, p<0.0001; Fig. 3b).

**Figure 3.**
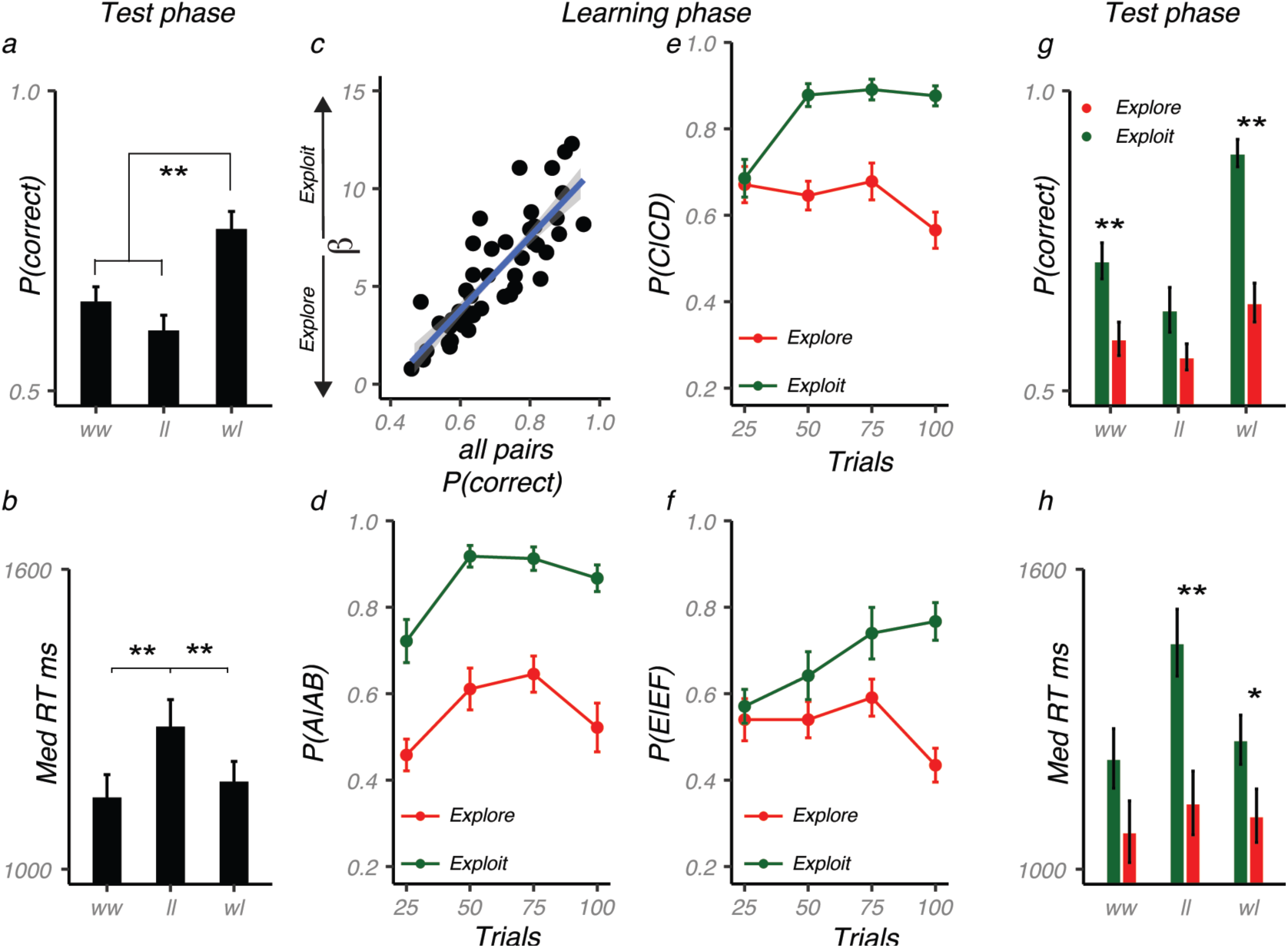
The exploit/explore trade-off in future decisions. Percentage of correct responses (a) and median reaction times (b) in the test-phase. c) Participants with an exploitative choice strategy learned well by mostly choosing the optimal options during the learning phase (d-f), and were more accurate (g) but slowed (h) in the test-phase; especially for the mostly neglected lose-lose pairs. The groups in plots d to h were created with a median split on β, and plotted to illustrate learning differences over time. Error bars represent SEM. **=p<0.01, *=p<0.05

To understand how the experience of conflict is influenced by the uncertainty associated with learned values that arises from information sampling, we quantified such sampling via the softmax β parameter estimated from the reinforcement learning model. Higher estimates of β index a greater tendency to exploit higher valued stimuli and as such predicted higher accuracies (r_AB_=0.79, r_CD_=0.74, r_EF_=0.47; all p’s<0.01; Fig. 3c) with steeper learning curves (Fig. 3d-f) in the learning phase. As noted above, however, we posited that such exploitation would increase the uncertainty about the values of under-sampled loss stimuli in future test-phase choices. A repeated measures ANOVA with the between subject variable Strategy (Exploit/Explore; defined as the continues variable β) and within subject factor Conflict (win-win, lose-lose, win-lose) revealed that exploitation during learning was related to improved accuracy (F(1,43)=72.1, p<0.0001; please see Fig. 3g for a visualization based on a median split on β), but also prolonged reaction times in the test phase (F(1,43)=9.0, p<0.01; Fig. 3h). Critically, these effects were qualified by an interaction between Strategy and Conflict (accuracy: F(2,86)=3.4, p=0.04; RT: F(2,86)=6.9, p<0.01), revealing especially large costs for lose-lose decisions in exploiters. In particular, compared to explorers, exploiters exhibited the most prominent RT cost for lose-lose (t(42)=3.5, p=0.001) and less so for win-lose (t(42)=2.1, p=0.04), and win-win (t(42)=1.7, p=0.09). Similarly, although they performed more accurately overall, exploiters showed significant gains in accuracy only for choices involving a win stimulus (win-win t(42)=3.2, p<0.01; win-lose t(42)=6.3, p<0.0001), and not for lose-lose choices (t(42)=1.8, p=0.08).

Hence, while it is unsurprising that overall, participants performing more accurately during training also do so at test, these exploitative participants were characterized by relatively selective RT costs for the lose-lose choices in the test phase. These costs are expected given that they had not sampled these stimuli as much and hence should exhibit larger uncertainty when choosing among them. We next considered whether such RT costs were mitigated by response inhibition, separately from choice strategy.

### Control and Conflict-induced slowing

While prolonged decision-times (RT’s) in the test-phase were related to exploitative choice strategies, we also were interested to assess the role of response inhibition independently of learning and uncertainty. Previous work has attributed conflict-induced slowing to the same STN mechanism associated with outright response inhibition^10,12^ via either dynamic modulation of decision thresholds and/or an initial delay that precedes the decision-process^4^. Therefore, we additionally examined an independent measure of inhibitory control efficiency in the stop-signal task termed the stop-signal reaction time (SSRT, Fig. 4a). We hypothesized that if conflict-induced slowing is simply associated with more overall response inhibition (or a fixed increase in decision threshold), then subjects engaging this mechanism would exhibit more inhibition and slower conflict-induced RTs. If, on the other hand, conflict-induced slowing involves a transient threshold increase that then collapses, then efficient response inhibition should relate to *less* conflict-induced slowing. Moreover, for exploiters, this release of a transient brake should particularly censor the tail of the RT distribution, which would otherwise have more density due to uncertainty in the evidence.

**Figure 4.**
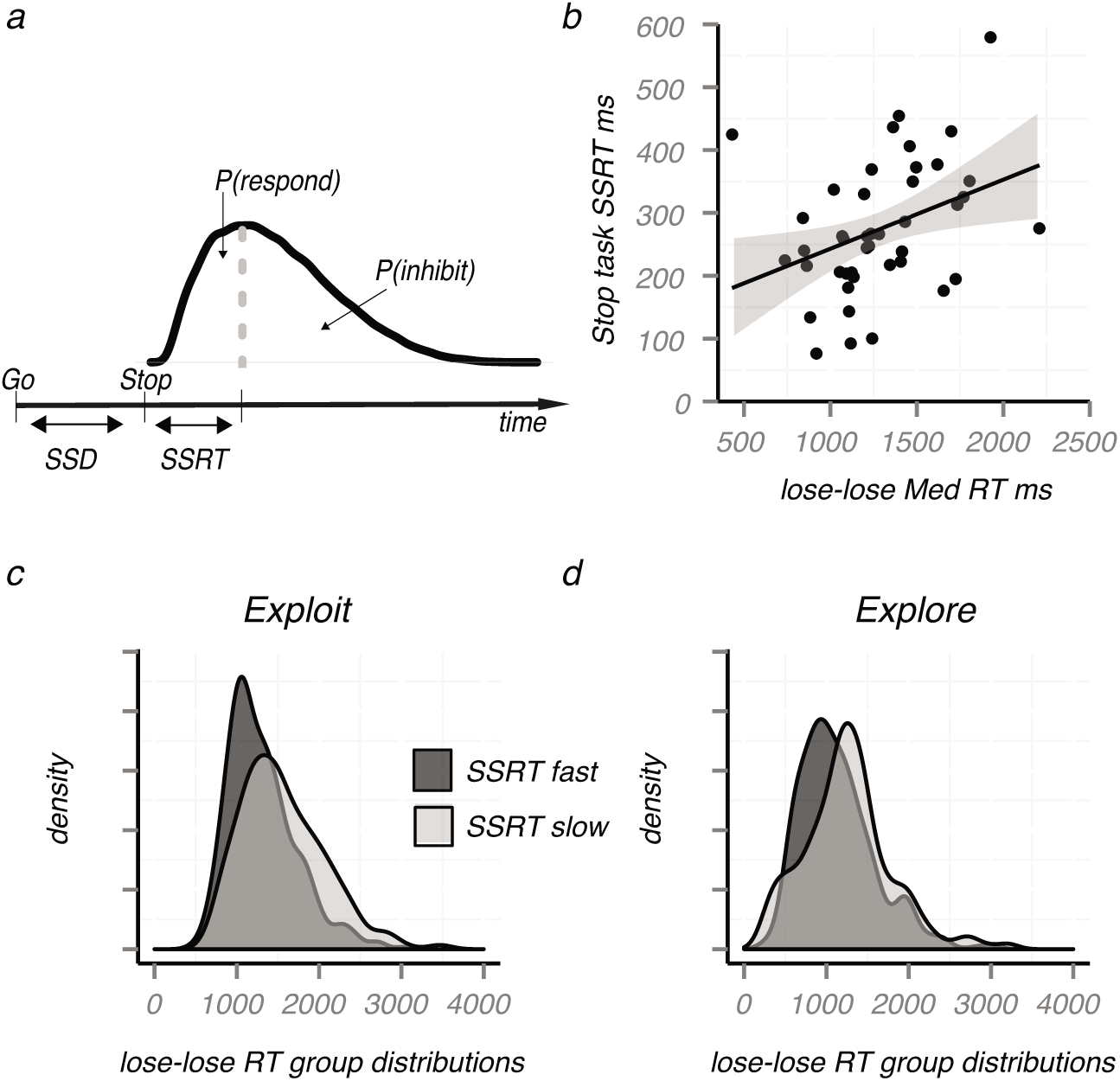
The efficiency to implement control predicts lose-lose slowing. a) Graphic representation of the race model estimation for SSRT. A distribution of go trial RTs is shown beneath the curve. SSRT represents the average time needed to suppress a planned response. The efficiency to stop (SSRT) predicted both response slowing and response times during lose-lose trials (b). Exploitative participants who were more efficient in inhibition showed a steeper decline in the tail of the lose-lose reaction time distribution (c), this was not seen for explorative participants (d). Median splits were used to create the Exploit/Explore or fast/slow SSRT groups.

Indeed, overall, faster SSRTs (more efficient inhibition) were related to faster lose-lose response times (r=0.36, p=0.02; Fig. 4b), and less slowing relative to low conflict win-lose trials (r=0.37, p=0.02). (No such relationship was seen for win-win RT; p=0.27). SSRT was unrelated to exploration vs exploitation in choices during learning (r=0.18, p=0.24), suggesting that the two factors might contribute independent variance to the lose-lose decision times. Indeed, a multiple regression showed a significant contribution of both β (b_β_=52.00, t(40)=3.356, p=0.002), and SSRT (b_ssrt_=0.92, t(40)=2.115, p=0.041) to lose-lose response times. Furthermore, while the inhibition effect was observable in both exploiters and explorers - consistent with an independent effect of SSRT on implementing and releasing the brake - its impact on the tail of the distributions was observed only in exploiters (Fig. 4c,d). This result is consistent with the notion that exploiters have more uncertainty about action outcomes, and hence without an efficient brake they exhibit longer tails. SSRT’s were not related to lose-lose accuracy performance (p=0.26).

These results explain lose-lose RT as a function of both choice strategies (previous sampling of information and hence uncertainty) and active but transient inhibitory control. Highly slowed participants were exploitative during learning, and inefficient in the implementation of a fast brake.

### The efficacy of control in the STN during full stops and conflict

At the neural level, the STN is well known for its role in global stopping (fast full brake) and the modulation of decision requirements. To evaluate how our behavioral observations relate to this literature, the time-course of activity within the STN was estimated for both the stop-signal task and the test-phase of the reinforcement learning task. Multiple regressions were then used to evaluate how the STN activity in each task relates to one’s efficacy to inhibit a planned response (SSRT), or choice strategy β.

In the stop-signal task, the estimated STN activity (Fig. 5a) was strongest for failed stop trials, corroborating a recent 7T study focusing on the STN in this task^28^, and possibly reflects a reactive engagement to correct for the failure to stop (we return to this result in the discussion). Notably, efficient inhibition, as indexed by SSRT, was marginally correlated with the estimated STN response *only* when participants succeeded to suppress a planned response on time (i.e., successful stop trials); such that higher early BOLD responses in the STN were related to faster or more efficient inhibition times (Fig. 5bc). As expected, no relationship was observed between the STN BOLD response and β, when participants were engaged in the stop-task.

**Figure 5.**
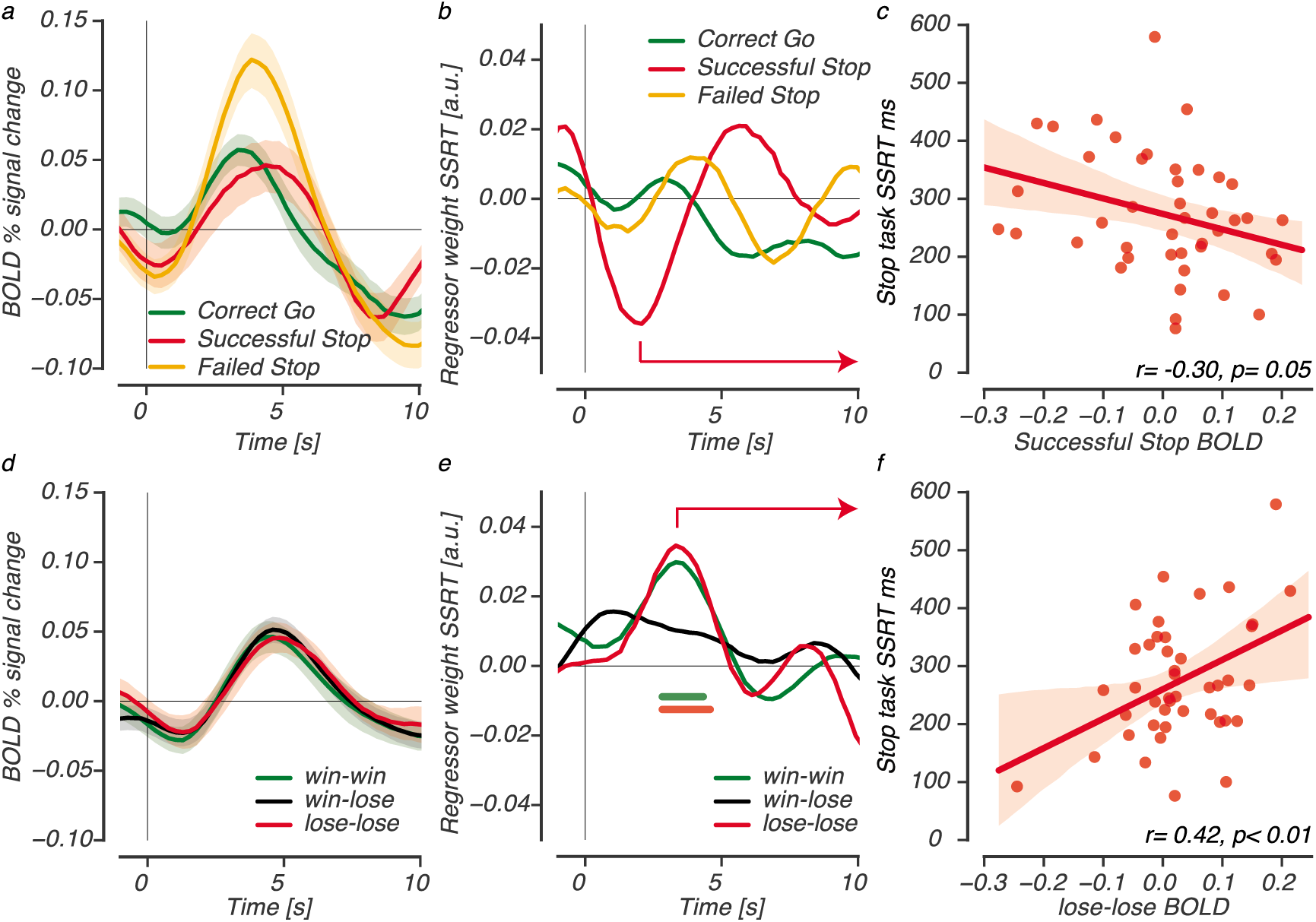
The efficiency to stop predicts the STN BOLD response in conflict and full response cancellation. The FIR-estimated STN BOLD signal time-course for trials in the stop-signal task (top panel) and the test-phase after learning (bottom panel), with the estimated regression coefficients SSRT shown for each trial type (mid panel). SSRT differentially related to activity patterns of the STN when inhibition was successful in the stop task (marginal effect with p=0.05), or with the experience of conflict in test-phase trials. The horizontal lines show the interval in which SSRT contributed significantly to the multiple regression, for the conflicted lose-lose (red) and win-win (green) trials. The right panel highlights the differential relationship across task, with drawn correlation plots for successful stop trials (c), and the slowed lose-lose trials after learning (f).

We then turned to the test-phase of the reinforcement learning task to understand how STN activity relates to the observed behavioral relationships between slowing and control. Overall, the estimated STN response was very similar across all trials of the test phase (Fig. 5d). The multiple regression, however, showed that STN response to high conflict (win-win and lose-lose), but not low conflict win-lose trials, was directly related to SSRT (Fig. 5e). STN activity was unrelated to β, pointing towards distinct effects of inhibitory control and choice uncertainty on response slowing. The positive relationship between SSRT and lose-lose STN activity (Fig. 5f) corresponds to the behavioral observation that more slowing was tied to longer SSRT’s, consistent with the notion that it results from the inefficient implementation of a transient STN brake.

### Conflict-induced slowing in cortico-basal ganglia pathways

Finally, we used the model-driven ancestral graphs approach (please see methods section for a detailed explanation) to analyze the information flow between PFC and BG during test-phase trials, and to explore how this interplay relates to the significant lose-lose slowing.

The evaluation of effective connectivity restricted the interplay between PFC and BG with the use of three pathways generally described in animal and human studies^20,29,30^ (Fig. 6a). Here, most projections terminate in the striatum (STR), from where two (out of three) pathways depart. A *direct*-*pathway* projects into thalamus via the globus pallidus interna (GPi) to facilitate action selection, while an *indirect*-*pathway* via the globus pallidus externa (GPe) can allow the integration of additional information by adaptively slowing the motor output. A third, *hyperdirect-pathway* directly projects from PFC into subthalamic nucleus (STN), and inhibits the thalamus output to primary motor cortex (M1) by exciting the GPi. These described PFC-BG pathways each play a specific (and therefore testable) role in the selection, regulation, or suppression of choices, and were therefore selected for the evaluation of seven potential connectivity networks to describe information flows, or connectivity, between the PFC and BG during test-phase decisions (please see methods section for the definition of all seven models).

**Figure 6.**
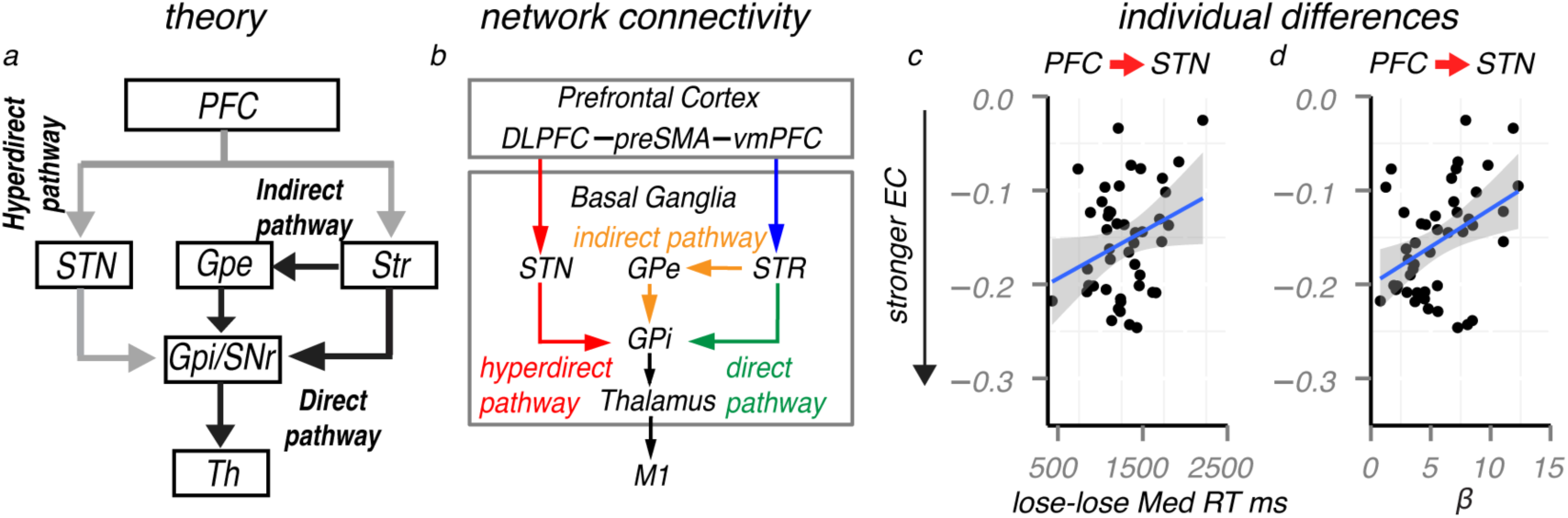
Schematic of the theoretical fronto-basal ganglia pathways and effective connectivity results. a) Theoretical fronto-basal ganglia model with the direct, indirect and hyperdirect pathways. Gray arrows represent excitatory connections; black arrows represent inhibitory connections. b) Graphical representation of the most representative effective connectivity network for all test-phase trials. Directed arrows represent effective connectivity (EC); undirected lines represent functional connectivity. For lose-lose trials, weaker PFC-into-STN connections related to prolonged response times (c) because of an exploitative choice strategy in the past and more uncertainty about the lose-lose options (d).

Concurrent with the literature, random effects analysis across the whole group indicated that a connectivity network comprising the direct, indirect and the hyperdirect pathway best describes the pattern of activity during all choice trials (Table 1). Fig. 6b shows the graphical outline of this model with functional connectivity (undirected relationship) between all PFC nodes (i.e., DLPFC, preSMA, vmPFC) and effective connectivity from each PFC region into the striatum (STR) and STN. Within BG, effective connectivity was defined from STN into GPi, Striatum into GPe, GPe into GPi, GPi into thalamus, and finally Thalamus into primary motor cortex (M1) to select a response. To better understand the top-down dynamics we next focused on connection strengths (regression values derived from ancestral graphs) from PFC into STN or STR in two steps.

**Table 1.** BIC values for model fits across test-phase conditions

| Model specification | win-win |  | lose-lose |  | win-lose l |  | win-lose m |  | win-lose s |  |
| --- | --- | --- | --- | --- | --- | --- | --- | --- | --- | --- |
|  | Left | Right | Left | Right | Left | Right | Left | Right | Left | Right |
| 1 PFC+BG direct | 18045 | 18222 | 18333 | 18453 | 12928 | 13321 | 13281 | 13306 | 13192 | 13226 |
| 2 PFC+BG indirect | 17491 | 17507 | 17731 | 17753 | 12535 | 12900 | 12862 | 12905 | 12743 | 12748 |
| 3 PFC+BG hyperdirect | 18709 | 18644 | 18987 | 19036 | 13367 | 13678 | 13674 | 13653 | 13636 | 13612 |
| 4 PFC+BG direct&hyperdirect | 17831 | 17906 | 18133 | 18320 | 12768 | 13119 | 13056 | 13082 | 13031 | 12986 |
| 5 PFC+BG direct&indirect | 17427 | 17453 | 17681 | 17705 | 12480 | 12832 | 12801 | 12855 | 12667 | 12673 |
| 6 PFC+BG indirect&hyperdirect | 16660 | 16614 | 16902 | 16918 | 11962 | 12296 | 12235 | 12279 | 12104 | 12116 |
| 7 PFC+BG direct&indirect&hyperdirect | 16588 | 16564 | 16848 | 16869 | 11906 | 12215 | 12176 | 12218 | 12018 | 12029 |

*Lower BIC values indicate a better balance between the variance and bias of the estimated model connections.*

First, the strength of top-down connections was investigated using a repeated measure ANOVA with the factors Conflict (win-win, lose-lose, win-lose) and Connection (PFC->STN, PFC->STR). There was a main effect of connection: PFC effective connectivity towards the STR (Mean=-0.17, SD=0.007) was stronger compared to that toward STN (Mean=-0.15, SD=0.007; F(1,44)=10.38, p=0.002). There were no additional main effects or interactions modulated by conflict.

Second, we explored how decision durations (RT’s) in the test-phase relate to the strength of connectivity (estimated regression strengths) from PFC into either STN or STR. For lose-lose trials, weaker, or disrupted, PFC-into-STN connectivity was related to longer RT’s (r=0.30, p=0.048; Fig. 6c) and more slowing (in comparison to the easy win-lose choices; r=0.32, p=0.03). These relationships were not observed for win-win or win-lose trials, or for PFC-into-STR connectivity (p’s>0.05). Because choice strategies and active control were both related to lose-lose RT (or slowing) in behavior, we next focused on the relationship between PFC-into-STN connectivity and β, or SSRT. The strength of PFC-into-STN connectivity correlated significantly with past choice strategies (r=41, p<0.01; Fig. 6d); such that participants with the most exploitative strategies, and hence most uncertain about values of lose-lose options, exhibited the weakest PFC-STN communication. Critically, a multiple regression of both β and reaction times on the PFC-into-STN connectivity (omnibus R^2^=0.18, F(2,42)=4.655, p=0.015), revealed that only β contributed significantly to this regression (b_β_=0.0068, t(42)=2.19 p=0.034), suggesting that the previously described relation to RT was mediated by uncertainty. In contrast, no relationship was found between SSRT and PFC-into-STN connectivity (p=0.18). The lack of a relationship between fronto-subthalamic connectivity and stop-task performance is consistent with our previous work focusing on the stop-signal task^11,24^, and possibly relates to the fast signal conduction within the hyperdirect pathway.

In sum, the directed PFC-into-STN connectivity correlated with the slowing of RT’s in lose-lose trials as a function of uncertainty, but not with the efficacy of inhibitory control implementation itself. These findings suggest that prolonged lose-lose decisions could result from: 1) weaker PFC-into-STN communications because of choice uncertainty, and 2) one’s efficacy to initiate and release a brake through the STN.

## Discussion

A large body of work focuses on the neural mechanisms of reinforcement learning and value-based decision making, and how animals and humans can optimize learning and choice performance in stochastic environments. However, here we provide evidence for a tradeoff: subjects that appear to perform better during learning are less able to quickly avoid the worst of low value options in a later generalization test. Because exploitative subjects during learning did not sample the less valuable options, they obtained less information about their precise probabilities. The concomitant increased choice uncertainty for later decisions and was marked by altered communication strengths from the PFC into STN and slower response times. We additionally focused on the mechanisms of this lose-lose slowing and the related neural response in the STN to show how both relate one’s ability to rapidly and transiently suppress all the preponent muscle buildup.

The fronto-subthalamic connections are thought to support conflict-induced slowing, and allow the integration of additional information by slowing or fully suppressing the motor output^2,4,12,15,17^. With the use of a model-driven connectivity approach, we found that the dynamic coupling between PFC and STN was weakest for the most uncertain and slowed participants. This relationship was further clarified by three additional findings. First, we observed that the magnitude of lose-lose slowing is best explained by considering not only past choice strategies (and hence uncertainty), but also, independently, stop signal reaction time (SSRT). The SSRT is an estimate for one’s efficiency to implement control^31,32^ and extensively related to the STN, which provides a fast and transient brake on all responses^1,9,10,23,33–40^. Accordingly, those subjects who were least efficient at rapid response inhibition (long SSRTs) exhibited more lose-lose slowing and a stronger STN surge during high conflict trials. Moreover, while no direct relationship was observed between choice strategies and SSRT, the fast (and transient) implementation of control was especially helpful in the prevention of overly slow lose-lose choices, especially for uncertain exploitative learners.

In the last decade two parallel lines of literature have focused on the specific role of the STN in the modulation of evidence requirements / decision threshold adjustments^2,4,17,41^, or full response suppression^9,35,36,42^. Response conflict has been consistently associated with the adaptation of evidence requirements^15,25,43,44^, including win-win and lose-lose conflict after reinforcement learning^2,3,45,46^. In this study, we evaluated the efficacy of control against the STN BOLD response during the experience of conflict, after learning, and in a separate task during full response suppression. We observed that the efficacy to implement a fast and full brake on all responses (SSRT) is differentially related to the STN in each process.

In the stop-signal task, the activity pattern of the STN was only related to stopping times when inhibition was successful. Here, the rise of the STN BOLD was highest for fast inhibitors. We note that this effect was only marginal in the 43 participants evaluated but consistent with the literature describing the STN in the stop task with rodents, or humans^9,10,38,47^. The strongest BOLD response in the STN was observed for trials where participants failed to inhibit a response on time^28,34^. Here, participants fail to inhibit the growth of activity for the go decision on time, and as a result might compensate by activating the STN without restraint or any regulation^48,49^. This compensation effort could increase estimates of the slow BOLD response, but as observed, should have no causal contribution to the stop process for which the average inhibition time is estimated with SSRT.

In contrast to successful stop trials, slower inhibition times were associated with increased STN BOLD responses in the evaluation of value-based decisions. Critically, however, this relationship was specific to the high conflict win-win and lose-lose trials, and not observed for easy win-lose decisions. Supporting the contrast between stopping and conflict, recent recordings from the STN have shown power increases in the STN to differ in the frequency range for conflict (2-8 Hz range)^2,50^, or response inhibition (13-30 Hz)^23,51,52^. Our results therefore suggest that a fast but transient STN brake, as posited by models showing a collapse in STN activity, might be helpful during conflict-choices.

The efficiency to suppress all responses correlated with the STN BOLD response during lose-lose and win-win conflict. Behaviorally, however, responses were only slowed and related to uncertainty, or SSRTs during lose-lose trials. Possibly, with the presentation of two negative options, the lack of information, negative value, and conflict all conspire to delay the selection process, or decision^4,53^. In contrast, when conflict is the result of two positive options (win-win) there is more information, and the STN activates to counterbalance only the most impulsive choices with the increase of evidence requirements^1,2,6^. The lack of response slowing for win-win choices can largely be attributed to the impact of predicted reward on the decision process itself. Indeed, when the normal counterbalancing function of the STN is disrupted win-win choices become even faster that the easy win-lose^1,4^.

The strength of fronto-STN connectivity or the magnitude of the STN BOLD both did not differ when compared between low conflict (win-lose), or high conflict (win-win, lose-lose) trials. Nevertheless, we observed selective relationships between uncertainty and fronto-STN connectivity for lose-lose slowing, or a relationship between control and the STN BOLD only at times of high conflict. These data imply fronto-subthalamic involvements and activity to be condition specific - despite any differences in magnitude. In the literature, condition specific relationships between PFC-theta and RT are found for learned high conflict choices – an effect that is reversed by deep brain stimulation of the STN – whereas no difference is found in overall theta power across conditions^2^. Moreover, the oscillatory activity of STN is related to behavior in opposing directions for low or high conflict conditions^17^. Our results complement these observations with the analysis of BOLD to show how high conflict responses can be specifically tied to disrupted fronto-STN dynamics, or inefficient control mechanisms.

Finally, previous time-sensitive reports have shown that the coherence between medial PFC and STN is increased in early periods of high-conflict^13,50^, with slower and more accurate responses when increases in the STN follow adaptations in medial PFC^22^. At first glance, our connectivity results contradict these findings. The critical difference here is that we evaluate connectivity, or co-activation patterns, with the use of trial-by-trial estimates of the slow BOLD response in PFC and BG nodes. In the STN, these BOLD estimates include both the rise (implementation), and fall (release)^4^ of the brake that is implemented to allow more time in conflicted decisions. With an identical rise, the trial-by-trial estimates of the STN BOLD should be lower with faster releases of the brake. We found stronger negative PFC-into-STN connections for the most certain participants, who responded faster, and chose the lesser options more often during learning. This pattern may suggest that when activity levels across PFC are raised sufficiently by information, the STN brake is released to allow choice. Consistently, PFC-into-STN connectivity was disrupted, and tied to slowing, for the most uncertain participants who mostly avoided the lesser options during learning. Future work should refine this interpretation with high temporal resolution approaches to evaluate connectivity in both early and late phases of conflict based decisions^54^.

To summarize, these results describe the profound lose-lose slowing as a function of past learning choices, and individual differences in active but transient response suppression through the STN (i.e., ‘hold and release your horses’). Moreover, they provide novel insights into the fronto-subthalamic (‘hyperdirect’) pathway involvement during the regulation of value-driven conflict.

## Methods

### Participants

All participants had normal or corrected-to-normal vision and provided written consent before the scanning session, in accordance with the declaration of Helsinki. The ethics committee of the University of Amsterdam approved the experiment, and all procedures were in accordance with relevant laws and institutional guidelines.

### Reinforcement learning task (RL-task)

The RL-task consisted of two phases; an initial reinforcement learning phase and a subsequent test-phase. During the learning phase, three different male or female face pairs (AB, CD, EF) were presented in random order, and participants learned to choose one of the two faces (Fig. 1a). Probabilistic feedback followed each choice to indicate ‘correct’ (happy smiley) or ‘incorrect’ (sad smiley). Choosing face-A lead to ‘correct’ on 80% of the trials, whereas face-B leads to ‘incorrect’. Other ratios for ‘correct’ were 70:30 (CD) and 60:40 (EF). Each trial started with a jitter interval of 0,500, 1000 or 1500ms to obtain an interpolated temporal resolution of 500ms. During this interval, a white fixation cross was presented and participants were asked to maintain fixation. Two faces were then presented left and right of the fixation-cross and remained on screen up to response. A white box surrounding the chosen face was then shown (300ms) and followed (interval 0-450ms) by feedback (500ms). Each trial had a fixed duration of 4000ms, and omissions were followed by the text ‘miss’ (2000ms). The test-phase contained the three face-pairs from the learning phase, and 12 novel combinations, in which participants had to select which item they thought had been more rewarding during learning. High conflict win-win trials were defined as choices that involved two previously rewarding stimuli (ie. AC, AE, CE), whereas high conflict lose-lose trials were defined as choices that involved two previously losing stimuli (BD, BF, DF). Low conflict win-lose stimuli served as controls for selection among novel pairs but which invoked little conflict (AD, AF, CB, etc). Test-phase trials (4000ms) were identical to the learning phase but no feedback was provided. In addition to the jitter used at the beginning of each trial, null trials (4000ms) were randomly interspersed during the learning (60 trials) and test (72 trials) phase. Across the whole task, each face was presented equally often on the left or right side, and choices were indicated with the right-hand index (left) or middle (right) finger. All stimuli were presented on a black-projection screen that was viewed via a mirror-system attached to the MRI head coil.

Before the MRI session, participants performed a complete learning phase to familiarize with the task (300 trials with different faces). In the MRI scanner, participants performed 2 learning blocks of 150 trials each (300 trials total; equal numbers of AB, CD and EF), and three test phase blocks of 120 trials each (360 total; 24 presentations of each pair).

### Stop-signal task

Each trial started with a white fixation cross followed by a male or female face stimulus indicating a left or right response with the index or middle finger of the right hand (Fig. 1b). Trials started with a random jitter interval of 0 to 1500ms (steps of 500 ms), during which a white fixation cross was presented in the center of the screen. A face stimulus was then presented for a period of 500 ms. On 30% of the trials, the go stimulus was followed by a high tone (stop signal). The stop signal delay (SSD) between the go stimulus and the stop signal was initially set at 250ms and adjusted according to standard staircase methods to ensure convergence to p(inhibit) = 0.5. For example, if a stop signal was presented and the participant responded (“failed stop”), then the SSD was reduced by 50ms on the subsequent stop trial; if the participant did not respond (i.e., “successful stop”), then SSD was increased by 50 ms. Instructions emphasized that participants should do their best to respond as quickly as possible while also doing their best to stop when an auditory stop signal occurred. Each trial had a fixed duration of 4000 ms, and trials were further separated by an occasional null trial with only a fixation (4000ms; 15 trials). Outside the scanner, participants performed a brief practice block of 30 trials to familiarize with the task. In the MRI scanner, participants subsequently performed a total of 150 trials (100 go trials, 50 stop trials). Faces used for the stop-task were not used in the RL-task.

### Reinforcement-learning model

We quantitatively characterized participants’ learning curves using a variant of the Q learning RL algorithm^55–57^, using hierarchical Bayesian parameter estimation, allowing us to separately estimate learning rates from choice stochasticity/exploration. Based on previous work we defined separate learning rate parameters for positive (α_gain_) and negative (α_loss_) reward prediction errors^56,58,59^. Q-learning assumes participants represent reward expectations for each stimulus/action (A-to-F). After observing a particular reward outcome, the expected value (Q) for selecting a stimulus *i* (A-to-F) on the next trial is updated as follows:

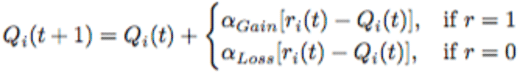

Where 0≤ α_gain_ or α_loss_ ≤1 represent learning rates, *t* is trial number, and *r=1* (positive feedback) or *r=0* (negative feedback). The probability of selecting one response over the other (i.e., A over B) is computed as:

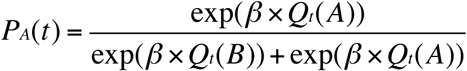

With 0≤β≤100 known as the inverse temperature governing the degree to which learned Q values are exploited. Higher estimates of β indicate that decisions are mostly determined by the relative difference in value (exploitation), whereas lower estimates show a more stochastic choice pattern but which facilitates better learning of the underlying values of the lesser options.

The Q-learning algorithm was fit to the learning-phase trials using a Bayesian hierarchical estimation method where parameters for individual subjects are drawn from a group-level distribution. This hierarchical structure is preferred for parameter estimation as it allows for the simultaneous estimation of both group level parameters and individual parameters, and confers greater statistical strength for estimating and recovering parameters^60–65^. Fig. 2a shows a graphical representation of the model. The quantities *r*_*i, t-1*_ (reward for participant *i* on trial *t*-1) and ch*_i,t_* (choice for participant *i* on trial *t*) are obtained directly from the data. The quantities α_Gi_, α_Li_ and *β*_*i*_ are deterministic, and are transformed during estimation by using their respective probit transformations Z’i (α'_Gi_, α'_Li_, β'_i_). The probit transform is the inverse cumulative distribution function of the normal distribution. The parameters *Z’i* lie on the probit scale covering the entire real line. Parameters *Z’i* were drawn from group-level normal distributions with mean *μ*_*z*_*’* and standard deviation *δ*_*z*_*’*. A normal prior was assigned to group-level means *μ*_*z*_*’* ∼ *N*(0,1), and a uniform prior to the group-level standard deviations *δ*_*z*_*’* ∼ *U*(1,1.5)^60,63^. Model fits were implemented in Stan^66,67^. Multiple chains were generated to ensure convergence, and evaluated with the Rhat statistics (i.e., all Rhats were close to 1.0)^68^. The right panel of Fig. 2 shows group-level posteriors on model parameters, and simulations from these parameters yield reasonable learning curves that match those observed empirically.

### Behavioral analysis

Accuracy rates and median reaction times (RT’s) were calculated for the RL-task test-phase and separated into high conflict win-win (ww; AC, AE, CE), high conflict lose-lose (ll; BD, BF, DF), and low conflict win-lose (wl; AD, AF, CB, CF, EB, ED) pairs, Fig. 1c. Pairs that were presented during the learning phase (AB, CD, EF) were excluded from the win-lose condition, so that all conditions only contained novel pairs. Repeated measures ANOVA’s with Tukey’s test were used to assess how conflict affects performance (RT and Accuracy). Conflict-induced slowing (or accuracy) was computed by subtracting median RT’s (or error rates) on win-lose trials from median RT’s (or error rates) on win-win or lose-lose trials. The stop-signal reaction time (SSRT) for the stop-task was estimated using the so-called ‘integration method’^31,43^. Pearson’s correlations, multiple regressions were used to focus on the relationship between conflict induced slowing/errors and choice strategies (β) or the efficiency to stop (SSRT), after reinforcement learning.

### Magnetic resonance imaging scanning procedure

The fMRI data for the RL-task was acquired in a single scanning session with two learning and three test-phase runs on a 3-T scanner (Philips Achieva TX, Andover, MA) using a 32-channel head coil. Each scanning run contained 340 functional T2*-weighted echoplanar images for the learning phase, and 290 T2*-weighted echoplanar images for the test phase (TR = 2000 ms; TE = 27.63 ms; FA = 76.1°; 3 mm slice thickness; 0.3 mm slice spacing; FOV = 240 × 121.8 × 240; 80 × 80 matrix; 37 slices, ascending slice order). After a short break of 10 minutes with no scanning, data collection was continued with a three-dimensional T1 scan for registration purposes (repetition time [TR] = 8.5080 ms; echo time [TE] = 3.95 ms; flip angle [FA]= 8°; 1 mm slice thickness; 0 mm slice spacing; field of view [FOV] = 240 × 220 × 188), and the fMRI data collection for the stop-task (335 T2* weighted echoplanar images; TR = 2000 ms; TE = 27.63 ms; FA = 76.1°; 3 mm slice thickness; 0.3 mm slice spacing; FOV = 240 × 121.8 × 240; 80 × 80 matrix; 37 slices, ascending slice order).

### Preprocessing

Preprocessing was performed using FEAT (FMRI Expert Analysis Tool) version 6.00, part of FSL (FMRIB's Software Library, www.fmrib.ox.ac.uk/fsl). The first six volumes were discarded to allow for T1 equilibrium effects. Preprocessing steps included motion correction, high-pass filtering in the temporal domain (σ = 50), and prewhitening^69^. All functional data sets were individually registered into 3D space using the participant’s individual high-resolution anatomical images. The individual 3D representation was then used to normalize the functional data into Montreal Neurological Institute (MNI) space by linear and non-linear scaling.

### fMRI analysis procedure and ROI selection

The analysis procedure of the fMRI data was twofold. First, an anatomically defined template of the STN region was used to explore how the STN BOLD response relates to SSRT (control) or β (choice uncertainty based on past learning) in the test phase after learning, and in the stop-signal task. The STN template was derived from a recent study using ultrahigh 7 tesla scanning^70^, and selected for its use in previous fMRI studies focusing on reinforcement based conflicted choices^13^, or response inhibition^24,25,28^.

Second, the model-driven ancestral graphs (AG) connectivity method was used for selecting the optimal network in describing PFC and BG co-activation patterns during test-phase trials (for the analysis of the stop-task using ancestral graphs please see^11,24,25^), and, to subsequently explore how the strength of PFC-BG connectivity in the optimal model relates to decision times, SSRT, or β. The regions of interest (ROIs) used for the AG analysis included: 1) masks based on a whole brain cluster-corrected analysis of the learning-phase fMRI data to identify regions that co-vary with trial-by-trial signed reward prediction errors, and 2) a priori selected PFC and BG anatomical masks for regions typically associated with fronto-BG decision-making, or choice evaluations^12,22,29,71,72^. Please see below for a detailed description of each step.

### Deconvolution analysis of the STN

To more precisely examine the time course of activations in the STN, we performed finite impulse response estimation (FIR) on the STN BOLD signals. After motion correction, temporal filtering and percent signal change conversion, data from the STN were averaged across voxels, and upsampled from 0.5 to 3 Hz. This allows the FIR fitting procedure to capitalize on the random timings (relative to TR onset) of the stimulus presentations and decisions in the experiment. For this analysis, stimulus onset was chosen as t0 of the FIR time course. FIR time courses for all trial types were then estimated simultaneously using a least-squares fit, as implemented in the FIRDeconvolution package^73^. Resulting single-participant response time-courses were then used to evaluate the contribution of SSRT and choice strategies for each timepoint separately, using multiple regression as implemented in the statsmodels package^74^. Here, alpha value for the contributions of SSRT and choice strategy was set to 0.0125 (i.e. a Bonferroni corrected value of 0.05 given the interval of interest between 0 and 8 s). Confidence intervals in Fig. 5 were estimated using bootstrap analysis across participants (n=1000), where the shaded region represents the standard error of the mean across participants (i.e. bootstrapped 68% confidence interval).

### Ancestral graphs method

To focus on fronto-basal ganglia dynamics when participants make reinforcement guided-decisions after learning, the fMRI data recorded during test-phase was analyzed using ancestral graphs^19^. Ancestral graphs (AG) infer functional or effective connectivity by taking into account the distribution of BOLD activation per ROI, across trials, per subject, and so are not dependent on the low temporal resolution of the time series in fMRI. A graphical model reflects the joint distribution of several neuronal systems with the assumption that for each individual the set of active regions is the same. The joint distribution (graphical model) of two nodes is estimated from the replications of *condition specific trials* (e.g., win-win, or lose-lose), and not from the time series. With this method, we can infer three types of connections: (i) effective connectivity (directed connection ->), (ii) functional connectivity (undirected connection -), and (iii) unobserved systems (bi-directed connection <->). Directed connections are regression parameters in the usual sense (denoted by *β*) and undirected connections are partial covariances (unscaled partial correlations; denoted by *λ*). The bi-directed connections refer to the covariance of the residuals from the regressions (denoted by *ω*). These three types of connections can be identified by modeling the covariance matrix (denoted by ∑) as:

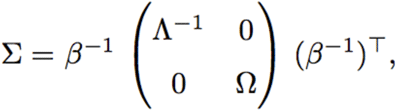

where *β* contains the regression coefficients, Λ contains the partial covariances, and Ω contains the covariances between residuals. A random effects model is used to combine models across subjects to then compare different models over the whole group using Bayes information criterion (BIC). The graph with the lowest BIC value will be selected.

To infer directions from the ancestral graph, it is required that a change in direction implies a change in probability distribution. This is not always the case. For example, a chain from A to B to C is in terms of conditional independencies equivalent to a chain with the directions reversed, that is from C to B to A (for more details see^19^). Two equivalent models, such as those just mentioned, will result in the same BIC value, indicating that directionality cannot be inferred. The most important structure is when two arrowheads meet (a collider). This will always result in a change in BIC value. The causal interpretations of the connections from an ancestral graph that is the best model according to the BIC can be briefly described as follows:

- A -> B: A is a cause of B [effective connectivity]
- A - B: A is a cause of B and/or B is a cause of A [functional connectivity]
- A <-> B: there is a latent common cause of A and B [missing region]

For a more detailed description and cautions on causal interpretations see^75^.

Individual (subject) fits are obtained by using an adjusted goodness-of-fit test, indicating whether the model explains the data well enough. To assess relative fit between the selected model and saturated model, the ancestral graphs method makes use of a modified version of the likelihood ratio (LR) test. For ancestral graphs, the modified LR test is defined as the ratio of the model of interest (hypothesis) and the unrestricted (saturated) model. The test is corrected for being overly sensitive because the data can deviate from normality slightly^76^.

The main differences between ancestral graphs (described in^19^) and dynamic causal modeling (DCM) or structural equation modeling (SEM) are: 1) inference is based on trial-by-trial variation in the estimated BOLD signal and not on the time series as in DCM or SEM because of the low frequency sampling in fMRI, 2) both functional and effective connectivity can be represented in a single ancestral graph, 3) a common unobserved (latent) cause of a connection can be detected, and 4) the definition of a circular system is only possible in undirected systems. The method of ancestral graphs relies on conditional independencies implied by the topology of the network. Therefore, different models (e.g., different directions of connections) result in different fits to the data. The differences between models is characterized by BIC, which combines both accurate descriptive (for the data at hand) and predictive (for future data) value.

### ROI definition for AG connectivity

AG connectivity was evaluated only for the test-phase trials of the RL task. The definition of ROI’s used for AG connectivity relied on 1) the analysis of the learning-phase fMRI data as to identify regions (voxels) within the striatum and vmPFC that correlate specifically with the signed reward prediction error (RPE), and 2) on previous work linking specific regions within the PFC and BG to value-driven decisions.

First, the two learning blocks were used to identify voxels within the vmPFC and striatum that respond to ongoing reward prediction errors during reinforcement-guided decision-making. For this purpose, the onset of each outcome was modeled as a separate delta function and convolved with the hemodynamic response function. We used a parametric GLM design with orthogonalized regressors where positive or negative outcomes were parametrically modulated by demeaned trial-wise prediction errors derived from the Q-learning model. Individual contrast images were computed for positive and negative error related responses and taken to a second-level random effect analysis using one-sample t-test. For the whole-brain analysis Z (Gaussianzied T/F) statistic images were thresholded using clusters determined by z > 2.3 (contrast positive RPE correlation) and a cluster-corrected significance threshold of p=0.05. Note, that this liberal threshold was only used for the definition of ROI masks that co-vary with RPE during learning; to be used only as masks for the evaluation of connectivity in the subsequent test-phase. During the test-phase feedback is no longer presented and internal representations of action values become vital to the selection process. Because reward prediction errors are thought to act as a teaching signal, ROI definition for the striatum [center of gravity (cog): 1, 5, -4] and vmPFC [cog: -3, 52, -1] nodes made use of the positive correlation RPE contrast in the learning phase, with the exclusion of voxels in the ventricles.

Second, a priori anatomical masks were defined for the following regions: preSMA [cog: (-) 9, 25, 50], DLPFC [cog: (-) 37, 37, 27], STN [cog: (L) -9, -14, -7; (R)10, -13, -7], globus pallidus interna (GPi) [cog: (L) -18, -8, -3; (R) 19, -7, -3], globus pallidus externa (GPe) [cog: (L) -19, -5, 0; (R) 20, -3, 0], thalamus [cog: (L) -10, -19, 7; (R) 11, -18, 7], and primary motor cortex (M1) [cog: -18, -26, 61]. All selected ROI’s were bilateral. The DLPFC template was obtained from a recent study, linking especially the posterior part to action execution^77^. The STN, GPe, and GPi templates were derived from a previous study using ultrahigh 7 tesla scanning^70^, thresholded to exclude the lowest 25% voxels, and then binarized. All other ROIs were created from cortical and subcortical structural atlases available in FSL.

### Single-trial parameter extraction for AG connectivity

For each ROI (anatomical or RPE based) we subsequently obtained a single parameter estimate (averaged normalized β estimate across voxels in each ROI mask) for each trial of the recorded test-phase, per subject. The average number of parameters (based on trials) per ROI was 71.1 (sd=1.7) for win-win, 71.4 (sd=1.3) for lose-lose, and 213.7 (sd=5.2) for win-lose. Misses were excluded from connectivity analysis. Connectivity analysis was conducted in R-Cran (version 3.0.2), including the packages ggm (version 1.995-3), graph (version 1.40.0), and RBGL (version 1.38.0).

### Model definition for AG connectivity

To examine how frontal and basal-ganglia nodes work together in selecting a response during the test phase, model fits were performed on the following trials: 1) win-win, 2) lose-lose, and 3) win-lose choices. A set of seven potential choice models containing the *direct* (PFC – Striatum –GPi –Thalamus–M1), *hyperdirect* (PFC – STN–GPi –Thalamus–M1), or *indirect* (PFC – Striatum –GPe – GPi –Thalamus–M1) PFC-BG pathways was tested to find the most optimal model in explaining the pattern of activation in the predefined regions. PFC consisted of vmPFC, DLPFC and preSMA, and each PFC region was defined to project into BG (see above for specification in the separate pathways). Because all PFC regions projected into BG,connections between PFC nodes could only be defined as undirected (functional connectivity). To optimize fits, all models were evaluated separately for left (right hand index finger) and right (right hand middle finger) responses (Table 1), and win-lose trials were first subdivided into three smaller chunks based on value-differences between pairs (small, 30; medium, 40; large, 50). Because connection strengths did not differ for win-lose divisions, parameter estimates of the winning model were averaged for the win-win, lose-lose, and win-lose condition to align with the behavioral analysis. To compare the contribution of each model with the BIC criterion all nine regions were always entered into the model, but the defined relationship (or connections) among regions varied across models.

## Data Availability

The code and processed files supporting the findings can be downloaded from: https://github.com/sarajahfari/Control_Conflict.git. The raw data is available from the corresponding author in BIDS format upon reasonable request.

## Acknowledgments

This work was supported with an ABC grant from the university of Amsterdam and National Science Foundation grant #1460604 to MJF.

## Author contributions

SJ, KRR and MJF designed the study. SJ collected the data. SJ, AC, TK, and LW contributed novel methods. SJ and TK analyzed the data. SJ and MJF wrote the manuscript.

